# Towards a standard approach to investigating the Thermal Load Sensitivity of photosystem II via chlorophyll fluorescence

**DOI:** 10.64898/2026.04.09.717599

**Authors:** Pieter A. Arnold, Rosalie J. Harris, Sabina M. Aitken, Max M. Hoek, Alicia M. Cook, Andy Leigh, Adrienne B. Nicotra

## Abstract

Evaluating the drivers of variation in plant thermal tolerance limits requires a clearer understanding of how methodological matters can lead to different tolerance estimates. Chlorophyll fluorometry – to measure the temperature-dependent change in *F*_V_/*F*_M_ – is a well-established approach to derive tolerance thresholds of photosystem II (PSII) in plants, but one-off, time-specific thermal exposures do not consider the fundamental dose-dependent effect of heat. The resurgent thermal death time (TDT) approach integrates both the temperature intensity and the exposure duration to derive time-based critical temperature thresholds and sensitivity parameters. We build upon this foundation to develop a protocol for evaluating thermal load sensitivity (TLS; non-lethal heat stress) of PSII in plants. Through five experiments across four diverse species, we tested the moderating effects of light, leaf sectioning, time since collection, and the temporal dynamics of *F*_V_/*F*_M_ recovery. There were dramatic changes in tolerance threshold estimates based on thermal load (i.e. dose-dependent) effects on *F*_V_/*F*_M_, and strong effects of light intensity during heat and the presence of light post-heat. We offer recommendations pertaining to method implementation and discuss future empirical avenues. Appraising cumulative heat stress will enhance the utility of thermal tolerance estimates – the TLS approach outlined here moves us toward a new standard.

## Introduction

Temperature is the primary determinant of species distribution and evolution (Nievola *et al*. 2017). Yet, we still struggle to define a species’ thermal niche (Moore *et al*. 2023), or predict how performance will be affected by climate warming and increasingly more frequent and intense extreme heat events (Seneviratne *et al*. 2012). It is increasingly clear that understanding the effect of thermal load (i.e. heat dose or cumulative heat stress) and species differences in sensitivity or capacity to avoid or repair heat-damage, is the critical next step for thermal ecology and ecophysiology (Jørgensen *et al*. 2021; Ørsted, Jørgensen & Overgaard 2022; Noble *et al*. 2026). A theoretical framework for modelling thermal load sensitivity across plants and animals has recently been developed (Arnold *et al*. 2025a). So too, have the importance of those insights for scaling to broader impacts of extreme heat events at community and landscape scales been demonstrated (Evans, Hu & Michaletz 2025; Ellis-Soto *et al*. 2026; Noble *et al*. 2026). Here we build on those recent theoretical advances and emerging empirical works to develop a standard method for assessing thermal load sensitivity of leaf tissue.

Research conducted over the last five decades has significantly advanced our understanding of plant heat stress (e.g. Christiansen 1978; Krishnan, Nguyen & Burke 1989; Porter, Nguyen & Burke 1994; Knight & Ackerly 2002; Hasanuzzaman *et al*. 2013; Geange *et al*. 2021; Jagadish, Way & Sharkey 2021). Heat tolerance is generally greater in species from warmer environments or with greater climate variability, but there is substantial unexplained among-species variance in thermal tolerance (O’Sullivan *et al*. 2017; Zhu *et al*. 2018; Lancaster & Humphreys 2020; Perez & Feeley 2020; Slot *et al*. 2021; Briceño *et al*. 2025). There is also extensive capacity for acclimation of critical temperatures as a function of growth environment (Briceño *et al*. 2025), recent history of thermal extremes (Andrew *et al*. 2023; Harris *et al*. 2024; Alvarez *et al*. 2025), and variation in tolerance depending on the overall plant health (Aitken *et al*. 2026). Methodological differences among studies are a substantial source of variation (Arnold *et al*. 2021; Perez *et al*. 2021a; Perez *et al*. 2025). In part these sources of variation reflect that conventional thermal stress measures (e.g. critical thermal limits) focus primarily on extreme temperature thresholds and as such do not incorporate cumulative impact of exposure on physiological function (Hochachka & Somero 2002; Rezende, Castañeda & Santos 2014; Rezende *et al*. 2020; Michaelsen, Fago & Bundgaard 2021; Ørsted, Jørgensen & Overgaard 2022; Cook *et al*. 2024; Faber, Ørsted & Ehlers 2024). Thus, while these are efficient for assessing snapshots of thermal tolerance variation among or within species at a moment in time, they are of more limited value for assessing effects of sub-lethal and fluctuating temperatures that plants encounter in real growing conditions. There is an emerging recognition that studies need to incorporate both the temperature intensity and duration of exposure to thermal stress (Williams *et al*. 2008; Huey *et al*. 2012; Huey & Kearney 2020; Arnold *et al*. 2025a), which for plants, generally focuses on the photosynthetic apparatus (Neuner & Buchner 2023; Faber, Ørsted & Ehlers 2024; Posch *et al*. 2025).

Leaf temperatures vary extensively in a given environment over short periods of time. Leaf morphology, growth architecture, plant water status, air temperature, vapour pressure deficit, shade, and windspeed all contribute to leaf temperature dynamics (Leigh *et al*. 2012; Rey-Sánchez *et al*. 2016; Blonder *et al*. 2020; Cook *et al*. 2021; Kearney & Leigh 2024; Manzi *et al*. 2025; Middleby *et al*. 2025). Species with traits that predispose them to higher leaf temperatures may also have higher thermal tolerance, illustrating the importance of leaf vs air temperature (Sastry, Guha & Barua 2017; Manzi *et al*. 2025). The capacity of plants to moderate leaf temperature via plasticity in morphology or via transpirational cooling is an emergent factor in context of heat exposure (Michaletz *et al*. 2015; Drake *et al*. 2018; Arnold *et al*. 2025b). Thus, air temperature is a poor surrogate for leaf temperature and a static leaf critical temperature reflects neither the thermal history of the leaf nor the potential for repeated excursions to relatively moderate temperatures to contribute to cumulative heat load (Leigh *et al*. 2017; Javad *et al*. 2025; Pottinger *et al*. 2025). Nevertheless, these effects can be captured in the context of the thermal tolerance landscape and modelled over realistic thermal regimes (Rezende, Castañeda & Santos 2014; Rezende *et al*. 2020; Cook *et al*. 2024; Aitken *et al*. 2026).

Here, we introduce a method designed to assess Thermal Load Sensitivity (TLS) of photosystem II (PSII) in photosynthetic tissues. The TLS approach builds on the established concept of Thermal Death Time (TDT), which incorporates both the temperature stress intensity and exposure duration of the tissue (Rezende, Castañeda & Santos 2014; Ørsted, Jørgensen & Overgaard 2022). TDT has been used to determine the critical thermal limits and sensitivity to thermal stress with a legacy in ectothermic animal physiology, but more recently is being applied to plants (Neuner & Buchner 2023; Cook *et al*. 2024; Faber, Ørsted & Ehlers 2024; Aitken *et al*. 2026). The TLS framework applies those concepts to evaluating non-lethal temperature thresholds, so that aspects of damage avoidance, loss of function, repair, and recovery can be examined directly as a function of both intensity and duration of heat exposure, under natural or realistic fluctuations (Rezende *et al*. 2020). The method described here aims to operationalise those principles for PSII in plants (Arnold *et al*. 2025a).

PSII is widely considered one of the most thermally sensitive components of the photosynthetic machinery, and it is the focus of much thermal tolerance work to date. Chlorophyll fluorometry can be used to quantitatively assess the maximum quantum yield of PSII, *F*_V_/*F*_M_ (Berry & Bjorkman 1980; Maxwell & Johnson 2000). Measures of relative change in *F*_V_/*F*_M_ provide a powerful tool for assessing immediate impacts of temperature on photosynthetic machinery. However, because these measures are so responsive, there are several aspects of measurement conditions that need to be accounted for. Both light intensity and timing of measurement of *F*_V_/*F*_M_ prior to, during, and after heat stress can influence estimates of PSII thermal tolerance (Havaux 1992; Curtis *et al*. 2014; Buchner *et al*. 2015; Szymańska *et al*. 2017; Ferrante & Mariani 2018; Nie *et al*. 2025). This is because light regulates plant signalling pathways, including those related to heat shock proteins and antioxidant defences (Li *et al*. 2021). Studies to date vary in light application (whether, when, and how much). In addition, the timing of *F*_V_/*F*_M_ measurement following thermal stress can vary from hours to days or weeks (e.g. Curtis *et al*. 2014; Haque *et al*. 2014; Nie *et al*. 2025). Further, thermal stress can be applied on entire or cut (discs or sections) leaves that differ in their excision time (Didion-Gency *et al*. 2025; Winter *et al*. 2025), leading to variable loss of function and damage and repair processes accumulating. For such assays to be readily comparable across experiments, standard protocols for setting light intensity levels and monitoring effects on *F*_V_/*F*_M_ after heat exposure are needed.

Our aim is to provide a resource for understanding and implementing the TLS method to *F*_V_/*F*_M_ in plants. Here we take into consideration the temperature intensity and exposure duration, as well as the issues around light level and timing of measurements outlined above. We aim to establish a robust tool to assess thermal tolerance of plant photosynthetic tissue under environmental stress, allowing for consistent comparisons across different experiments, species, and growth forms.

## Materials and Methods

### Species selection and experiment foundations

We evaluated the impact of temperature intensity and exposure duration, light conditions, leaf sectioning, and the time post-heat treatment on the proportional change of maximum quantum yield of PSII (i.e. photosynthetic efficiency, *F*_V_/*F*_M_) in four species, selected to represent diverse growth forms and leaf structures. Species were *Eucalyptus pauciflora* (Sieber ex Spreng), a tree with thick, long-lived leaves; *Lomandra longifolia* (Labill), a perennial rhizomatous rush; *Populus nigra* (L.), a deciduous cottonwood; and *Vicia faba* (L.), a bean commonly cultivated as a cover crop, all grown outdoors in ground in Canberra, ACT, Australia.

Drawing on the foundational work of Rezende, Castañeda and Santos (2014), Neuner and Buchner (2023), and Cook *et al*. (2024), we developed a comprehensive protocol tailored to examine PSII thermal load sensitivity (TLS) under a range of controlled environmental variables (**Fig. 1**). We conducted five experiments to specifically explore the effects of variation in 1) light exposure after heat stress, 2) light intensity during heat stress, 3) leaf integrity (cut vs whole), 4) time since sample collection and 5) the effects of time after the heat treatment on the proportional change in *F*_V_/*F*_M_. We measured *F*_V_/*F*_M_ before and after exposing photosynthetic tissue to a spectrum of heat stress conditions, then derived standard heat tolerance thresholds for 10%, 50%, and 90% (*T*_10_, *T*_50_, *T*_90_) reductions from an initial *F*_V_/*F*_M_ value (Cook *et al*. 2024) and the temperature at which *F*_V_/*F*_M_ declines to an absolute value of 0.3 (*T*_0.3_), which is an assumed value for irreversible loss of function that is independent of the initial *F*_V_/*F*_M_ (Curtis *et al*. 2014; Aitken *et al*. 2026). Various other thresholds for the temperature at initial *F*_V_/*F*_M_ decline (e.g. ‘*T*_crit_’, *T*_5_, *T*_15_) and the point of significant loss of function (e.g. *T*_95_) have been used previously and often chosen arbitrarily as a point of comparison (Perez *et al*. 2021b; Tiwari *et al*. 2021; Kunert & Hajek 2022; Valliere, Nelson & Martinez 2023; Posch *et al*. 2025; Winter *et al*. 2025).

**Figure 1.**
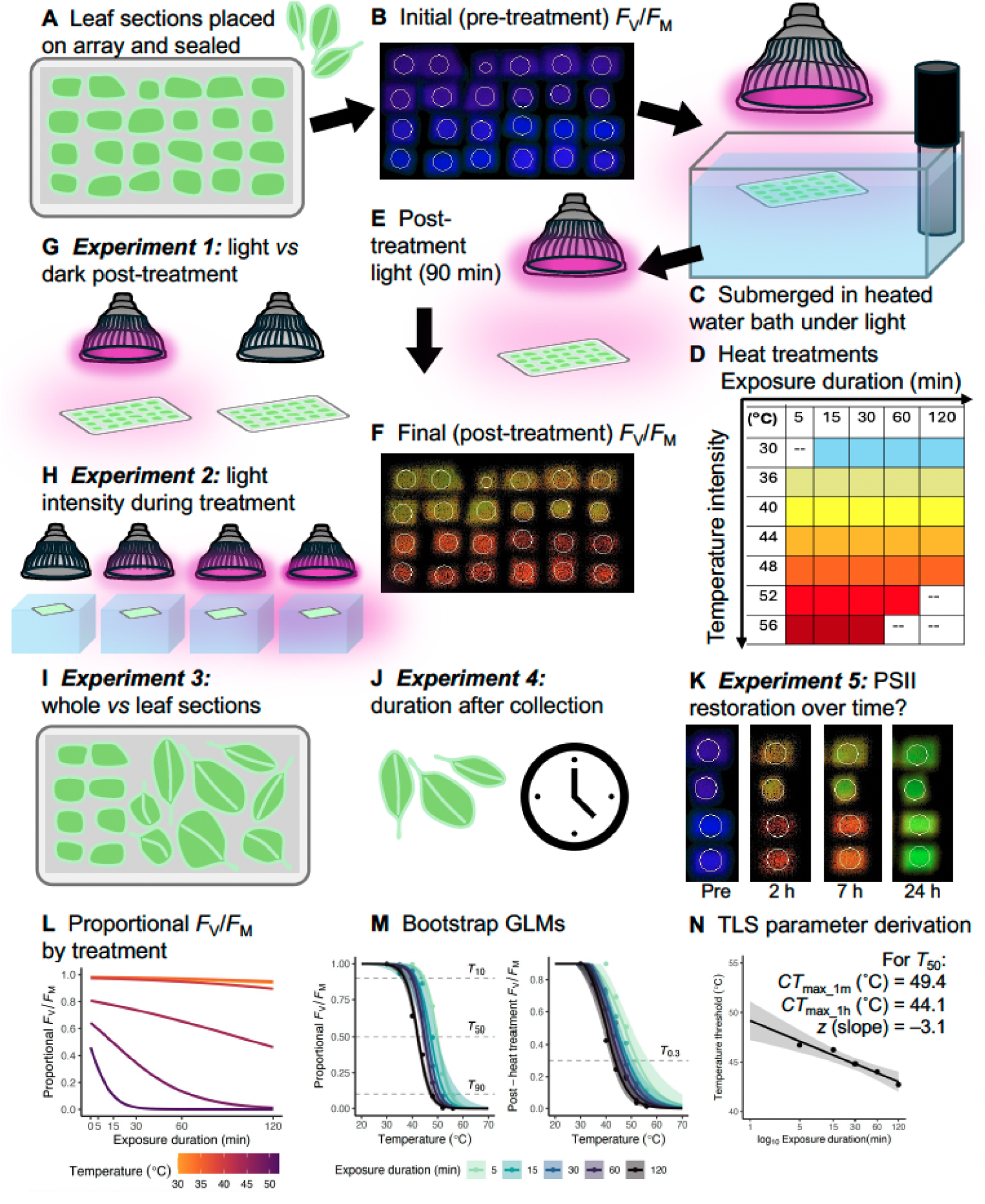
The general protocol for thermal load sensitivity (TLS) of PSII using *F*_V_/*F*_M_, the five experiments we used to explore the moderating effect of light and other experimental factors, and the model fitting procedure. **A** Whole leaves collected from the field were cut into subsections and placed on a paper array then sealed in a waterproof plastic sleeve. **B** Arrays were dark adapted and then pre-treatment *F*_V_/*F*_M_ was measured. **C** Each sealed array was exposed to 15 min of light prior to being submerged in a heated water bath under grow lights for **D** one of up to 31 different heat treatments (temperature intensity × exposure duration combinations). **E** After heat treatment, the arrays were placed under moderate light out of the water bath for 90 min. **F** Arrays were left in darkness overnight then post-treatment *F*_V_/*F*_M_ was measured. **G** Experiment 1 modified the light post-treatment. **H** Experiment 2 modified the light during treatment. **I** Experiment 3 compared whole vs leaf sections. **J** Experiment 4 explored timing of leaf collection. **K** Experiment 5 explored the temporal dynamics of restoration of PSII function. **L** Proportional *F*_V_/*F*_M_ compared across treatments. **M** Bootstrapped logistic GLMs for proportional and post-heat treatment *F*_V_/*F*_M_ allows derivation of threshold metrics (left: *T*_10_, *T*_50_, *T*_90_; right: *T*_0.3_) at each exposure duration. **N**. TLS parameters (*CT*_max_1m_, *CT*_max_1h_, *z*) then derived from a log_10_-linear model of the threshold metrics regressed against exposure duration.

### Temperature treatments and fluorescence assays

Unless otherwise noted, the following protocol for collection and assays was applied for each experiment. Short branches were cut from plants growing outdoors in the morning and immediately stored in sealed plastic bags with moistened paper towel. Leaf material was prepared as soon as possible by cutting ∼1 cm^2^ leaf sections, which were attached to an adhesive gridded paper array. Once leaf sections were arranged on arrays, they were enclosed in translucent plastic sleeves with a small volume of air (**Fig. 1A**) using a heat sealer (FoodSaver VS2198, Newell Brands, Atlanta GA USA). Prior to heat treatments, arrays of leaf samples were dark-adapted for 20 mins, followed by an initial *F*_V_/*F*_M_ measurement (**Fig. 1B**) using a Maxi-Imaging-PAM (Heinz Walz GmbH, Effeltrich, Germany). After this baseline assessment, the arrays were exposed to sub-saturating light of ∼700 µmol m^−2^ s^−1^ for 15 mins to light-adapt (Curtis *et al*. 2014).

Arrays of leaf sections were then completely submerged in temperature-controlled water baths (**Fig. 1C**). The temperature was maintained to within ± 0.2°C of the desired temperature using SousVide Precision Cookers (Kogan, Melbourne VIC Australia). Water bath temperatures were continuously monitored with type-T thermocouples (Omega Engineering, Singapore) connected to a datalogger (DataTaker DT85, Lontek, Glenbrook NSW Australia) to ensure temperature stability. Arrays were submerged ∼ 20 mm below the waterline of the water bath with a slim wire rack to ensue each array remained flat and parallel to the light source throughout the treatment. We suspended grow lights (15 × 1 W LEDs: 6 × 660 nm, 5 × 630 nm, 1 × 710 nm, 2 × 460 nm, 1 × White 3400 K) above each water bath and adjusted heights to ensure consistency in light intensity across arrays in the water baths by measuring photosynthetic photon flux density with a full spectrum quantum meter (MQ-500, Apogee Instruments, Logan, UT, USA). The heat treatment assays were conducted at ∼700 µmol m^−2^ s^−1^, which was considered moderate and sub-saturating, unless otherwise stated. Plants were exposed to temperatures between 30–56°C and across durations that ranged 5–120 mins (**Fig. 1D**). Not all combinations were assessed and these differed among experiments (e.g. only short to moderate durations for the highest temperatures and moderate to longer durations for the lower temperatures). After the heat treatment, arrays were removed from the water baths and the sealed plastic sleeves were opened to allow gas exchange. Arrays were put under the same moderate, sub-saturating ∼700 µmol m^−2^ s^−1^ as during the heat treatment at ambient temperature (22°C) for 90 mins post-heat treatment (**Fig. 1E**). Then, arrays were moved into darkness at ambient temperature overnight and a final measurement of *F*_V_/*F*_M_ was taken between 16–24 h later (**Fig. 1F**).

### Experiment 1: Light vs dark conditions post-heat treatment

We investigated whether light treatment after heat exposure affected the changes in *F*_V_/*F*_M_ because we expect a plant would naturally be exposed to light for several hours following a heatwave or a hot part of the day. Post-heat treatment, pairs of arrays from a given treatment were divided into two groups: one group was placed under light (700 µmol m^−2^ s^−1^) post-heat treatment as described above while the other group was put in darkness immediately after heat treatment until *F*_V_/*F*_M_ was taken 16–24 h later (**Fig. 1G**).

### Experiment 2: Effects of light intensity during heat treatment

Here we aimed to elucidate the effects of varying light intensities during heat treatment on the final *F*_V_/*F*_M_ of plants, recognising that heat stress typically occurs in conjunction with sunlight in natural settings. This involved subjecting leaf arrays to a range of light intensities (**Fig. 1H**) – 0 µmol m^−2^ s^−1^ (none, i.e. dark), 350 µmol m^−2^ s^−1^ (low), 700 µmol m^−2^ s^−1^ (moderate), and 1100 µmol m^−2^ s^−1^ (high) – while exposing them to temperatures between 30– 56°C for varying durations. *F*_V_/*F*_M_ measurements were taken before the heat treatment and 16–24 h later, with post-heat treatment as the light conditions per Experiment 1). We expected that sub-saturating light levels might induce protective mechanisms and offset heat effects, whereas higher light levels might exacerbate the heat effects, leading to more pronounced declines in *F*_V_/*F*_M_.

### Experiment 3: Comparison of whole leaves vs leaf sections

Using excised leaf sections or discs enables higher throughput in these assays. We therefore tested whether cutting the leaf altered stress responses relative to entire leaves, inclusive of the petiole (**Fig. 1I**). We subjected the whole leaves or leaf sections to a subset of temperatures (30, 40, 48°C) for a subset of durations (5, 30, 120 mins) used in the assays in Experiments 1 and 2, and measured *F*_V_/*F*_M_ as described above.

### Experiment 4: Influence of duration after collection on leaf responses

We next investigated how the time elapsed from collecting the leaves in the field to then using them in the laboratory affected the responses of the leaf tissue. We collected leaves at various intervals – immediately before experimentation, or 3 h, 12 h, or 24 h – prior to use in the experiment to assess how pre-experimental sample storage time affected heat stress (**Fig. 1J**). Using samples immediately following collection is often impractical, therefore understanding the viability of storing samples is important. Short branches or shoots were cut and stored in sealed bags with moistened paper towel for the various intervals until preparation. All samples were cut and assayed across the same subsets of temperatures and durations as in Experiment 3.

### Experiment 5: Effects of time post-treatment on restoring PSII function

Finally, to investigate the time course of thermal stress response and the potential for either recovery of function or lags in loss of function, we measured *F*_V_/*F*_M_ at varying intervals of 30 mins, 2 h, 7 h, 16 h, and 24 h after heat stress (**Fig. 1K**). This was done on a subset of samples from Experiment 3 above using leaf discs only.

### Statistical analyses

All analyses were conducted in R 4.5.1 (R Core Team 2023), using the additional packages *tidyverse* (Wickham *et al*. 2019) for data formatting and plotting, *lmerTest* and *lme4* (Bates *et al*. 2015; Kuznetsova, Brockhoff & Christensen 2017) for fitting models, and *emmeans* (Lenth 2023) for post-hoc tests.

For Experiments 1 and 2, we scaled post-heat treatment *F*_V_/*F*_M_ as a proportion of the initial (pre-treatment) *F*_V_/*F*_M_ value. A proportional *F*_V_/*F*_M_ value of 1 therefore indicates no damage (or functional impairment) to PSII from heat treatment while 0 indicates complete loss of function, and 0.5 represents 50% reduction from initial function (**Fig. 1L**). We then fitted logistic regressions using generalised linear models (GLM) including the temperature intensity × exposure duration treatments and their interaction with each respective experimental variable (e.g. for Experiment 1 the experimental variable was light) as predictor variables for the response variable of proportional *F*_V_/*F*_M_. we bootstrapped 1000 simulations with replacement to estimate uncertainty for GLM model predictions of proportional *F*_V_/*F*_M_ (Cook *et al*. 2024). The threshold values for loss of function (*T*_10_, *T*_50_, *T*_90_) were then estimated by identifying the temperature at which the predicted proportional *F*_V_/*F*_M_ reached 0.9, 0.5, and 0.1, respectively (**Fig. 1M**). We next fitted simple linear models onto the derived temperature thresholds from the bootstrapped GLM for each duration (**Fig. 1N**). From that linear model, we extracted the theoretical heat tolerance limit from 1-min exposure (*CT*_max_1m_, the *y*-intercept), a more biologically relevant heat tolerance limit from 1 h exposure (*CT*_max_1h_), and a thermal sensitivity parameter (*z*, the slope of the linear regression that describes the change in tolerance for 10-fold increase in exposure duration) (Rezende, Castañeda & Santos 2014). Following Aitken *et al*. (2026), the temperature at which final *F*_V_/*F*_M_ = 0.3 (*T*_0.3_) was determined from logistic models fitted to each duration treatment as above. The value of 0.3 was selected as a threshold at which significant irreversible damage has occurred to PSII (Curtis *et al*. 2014); it is a possible indication of irreversible damage that is not influenced by initial *F*_V_/*F*_M_ prior to heat treatment. These thresholds are expected to be ordered *T*_10_, *T*_0.3_, *T*_50_, *T*_90_ in ascending order of functional impairment, where *T*_0.3_ would be equivalent to *T*_50_ when a leaf has an initial *F*_V_/*F*_M_ value of 0.6.

Experiment 1 aimed to assess the influence of light conditions following heat treatment on the heat tolerance limits derived from proportional change in *F*_V_/*F*_M_. We fitted a linear mixed-effects regression (LMER) model with thresholds (*T*_10_, *T*_0.3_, *T*_50_, *T*_90_) as the response variable with light treatment (light vs dark), species, exposure duration, and all their interactions as fixed effects. The plant sample was included as a random effect to account for temporal and individual variation among sampling efforts. We then fitted LMER models on the TLS metrics (*CT*_max_1m_, *CT*_max_1h_, *z*) to primarily evaluate the effect of light treatment, but also differences among species, the threshold from which the TLS metrics were derived, and their interactions as fixed effects with sample as a random effect. We report ANOVA from each of these LMER models for this and all other experiments. Experiment 2 aimed to assess how light intensity during heat exposure affected the heat tolerance limits derived from proportional change in *F*_V_/*F*_M_. LMER models were fitted as above but with light intensity (0, 350, 700, 1100 µmol m^−2^ s^−1^) replacing light treatment (light vs dark).

Experiments 3, 4, and 5 were analysed with the same model structure. These experiments were smaller tests (fewer combinations of temperature intensity and exposure duration) of factors that could alter *F*_V_/*F*_M_, so we analyse the *F*_V_/*F*_M_ response rather than fitting models to derived thresholds and TLS metrics. Experiment 3 aimed to determine whether the physical state of the leaf (whole or cut into sections) affected the *F*_V_/*F*_M_ before and after heat treatment. We fitted a LMER with leaf integrity (whole vs cut), temperature and duration treatments (as a combined factor with 9 levels because their isolated effects were not the focus of these experiments), species, and their interactions as fixed factors, with sample as a random effect as above. The same model was fitted for Experiment 4, replacing leaf integrity with time since collection. For Experiment 5, we fitted a similar LMER with the time point of measurement replacing time since collection, on which we conducted pairwise post-hoc contrasts on estimated marginal means with Kenward-Roger degrees of freedom. This approach revealed whether *F*_V_/*F*_M_ changed from 30 min and 24 h post-heat treatment measurements in each temperature and duration treatment within species.

## Results

### Overview

Our work confirms that temperature intensity and exposure duration had cumulative effects on *F*_V_/*F*_M_ as expected: longer exposure duration and higher temperature significantly reduced *F*_V_/*F*_M_. Heat tolerance thresholds (*T*_10_, *T*_0.3_, *T*_50_, *T*_90_) declined significantly with exposure duration with few exceptions, and these thresholds have robust linear relationships with log_10_-transformed exposure duration that enables consistent prediction of TLS metrics (*CT*_max_1m_, *CT*_max_1h_, *z*). An understanding of the measurement conditions that can moderate these parameters is crucial and can be summarised as follows. 1) Light following heat stress can alter apparent heat tolerance, but sensitivity is similar; 2) light intensity during heat stress reduces heat tolerance; 3) cut leaf sections respond similarly to whole leaves; 4) sampled leaves can be kept for at least up to 24 h without confounding effects on *F*_V_/*F*_M_; and 5) recovery of *F*_V_/*F*_M_ function within 24 h is possible but the magnitude thereof is species-specific.

### Experiment 1: Exposure to light following heat stress can alter heat responses depending on species and thresholds

The four species differed strongly in their capacity to tolerate heat, with *T*_50_ thresholds ranging between 37–54°C. Exposure duration was consistently a significant factor in affecting heat tolerance thresholds (**Fig. 2; Table S1**). Light had a significant interaction with exposure duration for all thresholds, while light responses interacted with species for *T*_90_ (**Fig. 2; Table S1**). The various thresholds on the *F*_V_/*F*_M_*-T* curve (*T*_10_, *T*_0.3_, *T*_50_, *T*_90_) are affected by duration and light in a similar manner among all species except for *L. longifolia* (**Fig. 2**). The near-zero *z* (i.e. no temporal dependence) for *T*_50_ and *T*_90_ estimates in *L. longifolia* indicate that exposure duration was less important than temperature intensity in determining heat tolerance. The derived TLS metrics were all significantly lower when exposed to light (**Fig. 3; Table S2**).

**Figure 2.**
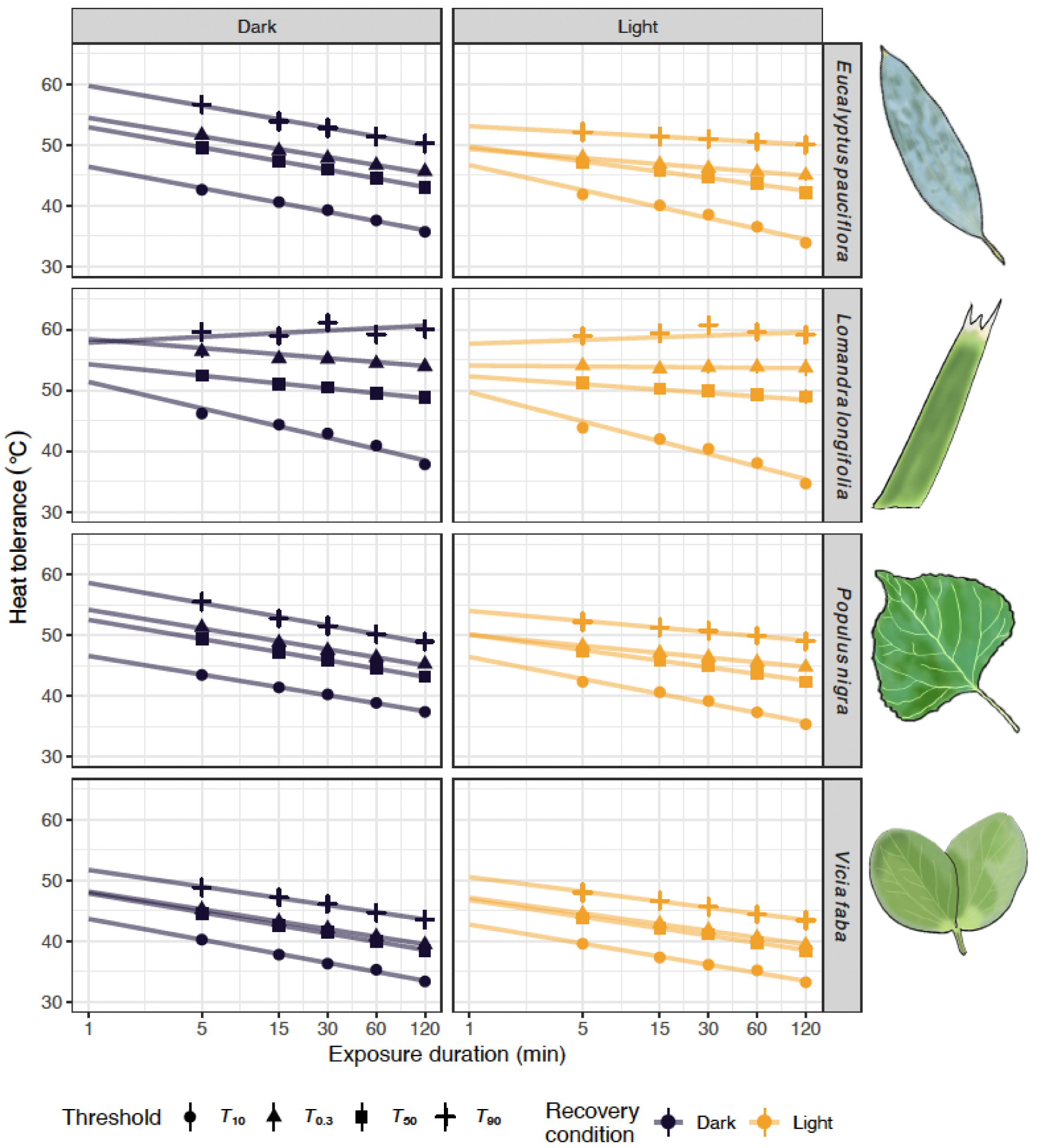
Experiment 1 findings: effects of post-treatment light conditions on heat tolerance and derived TLS metrics. Species and light recovery condition are separated by facets: 90 minutes of exposure to darkness (left, black) or moderate light (right, orange) after heat treatment are shown as left and right columns respectively, where each heat tolerance threshold (*T*_10_, *T*_0.3_, *T*_50_, *T*_90_) is shown by a different symbol. TLS metrics can be inferred from these fitted regressions such that *CT*_max_1h_ for a given threshold is the y-value at x=60, *CT*_max_1m_ for a given threshold is the y-value at x=1 (intercept), and *z* for a given threshold is the slope of the regression (plotted in Fig. 3). Note that the *x*-axis is on a log_10_ scale. Graphic of typical species’ leaves (not to scale) by P. A. Arnold.

**Figure 3.**
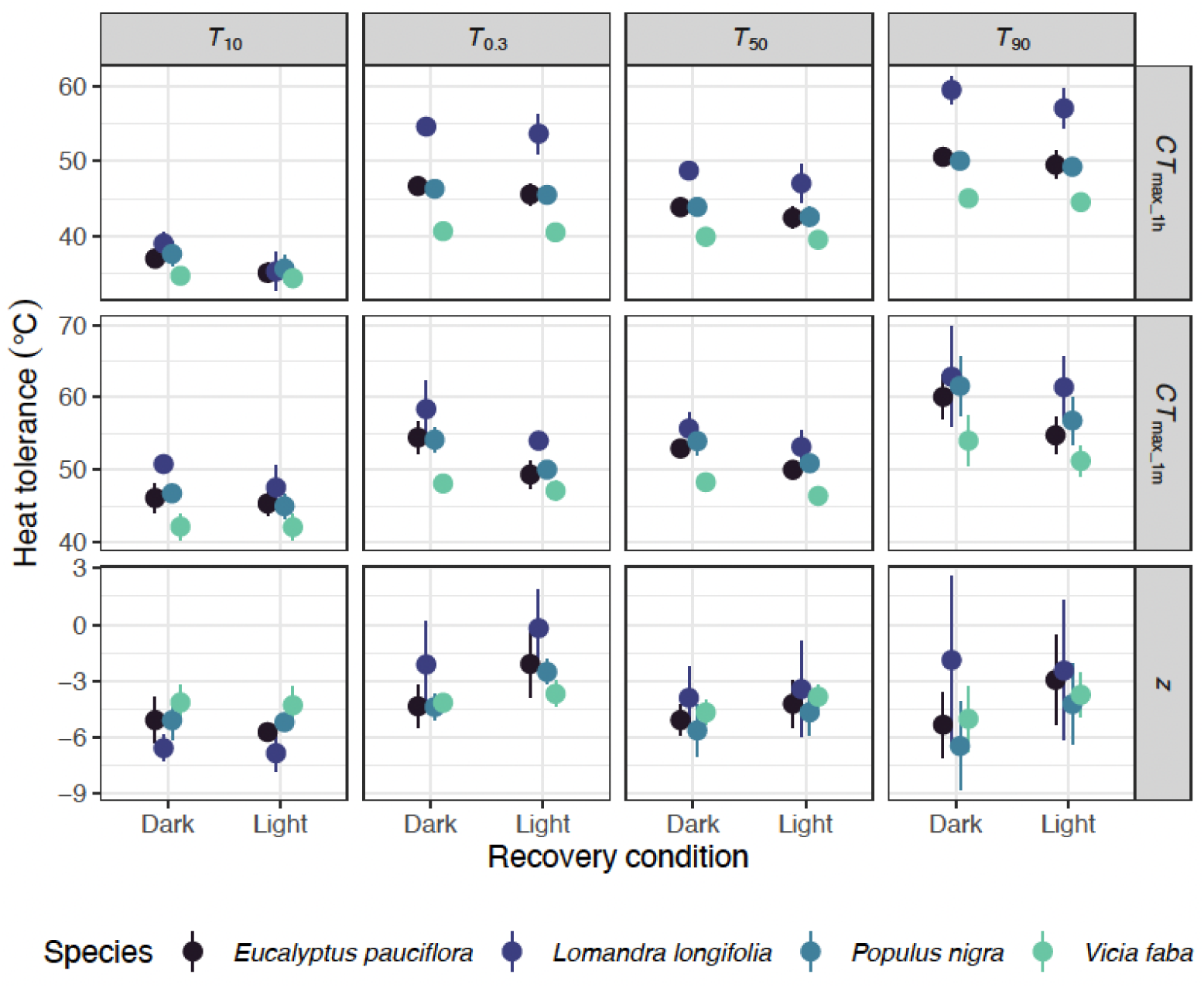
Experiment 1 findings: effects of post-treatment light conditions derived TLS metrics. Threshold (*T*_10_, *T*_0.3_, *T*_50_, *T*_90_) and TLS metrics (*CT*_max_1h_, *CT*_max_1m_, *z*) are each shown on separate panels. Species are shown by different colours and recovery condition is on the x-axis. Points and error bars are means ± SE.

### Experiment 2: Light intensity during heat reduces heat tolerance

To determine whether light exposure during heat stress exacerbated or ameliorated the cumulative heat stress we conducted further assays at four light intensities ranging 0–1100 µmol m^−2^ s^−1^. We found that light during heat stress significantly reduced the apparent heat tolerance, and this was consistent across each species and threshold (**Fig. 4; Table S3**). The higher the light intensity during heat stress, the lower the heat tolerance values (**Fig. 4**). This holds for all four species, which indicates that light intensity exacerbates rather than ameliorates the accumulation of heat damage (**Fig. 4**). The TLS parameters show that increasing light intensity lowers the derived *CT*_max_1h_ and *CT*_max_1m_ values, while *z* tends toward less negative values (**Fig. 4, Table S4**).

**Figure 4.**
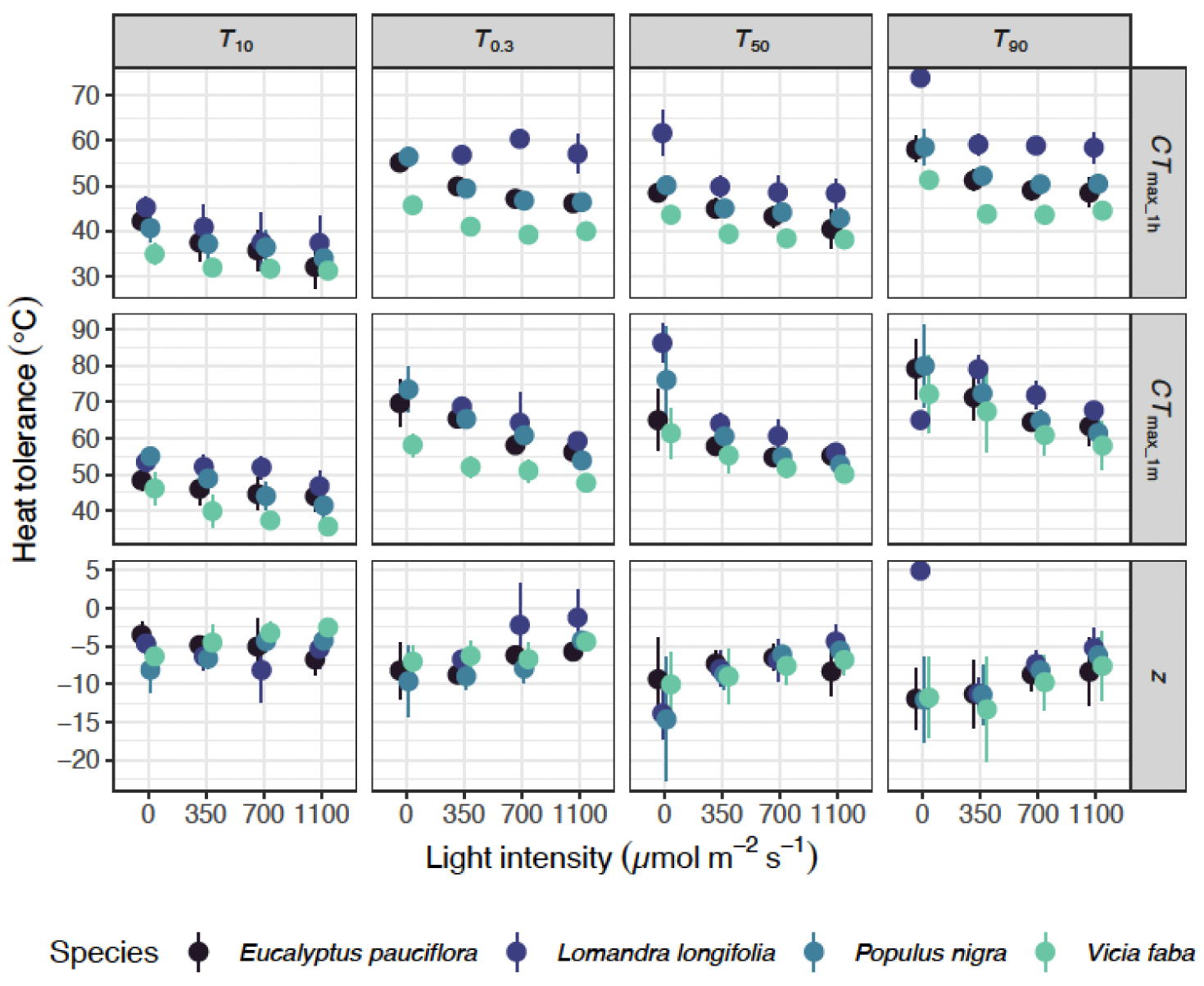
Experiment 2 findings: effects of light intensity during heat treatment on TLS metrics. Threshold (*T*_10_, *T*_50_, *T*_90_, *T*_0.3_) and TLS metrics (*CT*_max_1h_, *CT*_max_1m_, *z*) are shown in separate panels. Species are differentiated by different colours, and light intensity is on the x-axis. Points and error bars are means ± SE.

### Experiment 3: Cutting leaves into sections does not compromise the accuracy of heat tolerance measurements

To determine whether cutting leaves into sections altered the responses relative to whole leaves, we assessed the decline in *F*_V_/*F*_M_ at a subset of temperatures and exposure durations. Initial *F*_V_/*F*_M_ values were not affected by cutting the leaves: both cut and whole leaves had mean initial *F*_V_/*F*_M_ > 0.8. There was also no significant difference between cut and whole leaves for final *F*_V_/*F*_M_ (**Fig. 5**). As expected from Experiments 1 and 2, the temperature × duration combination and species had significant effects on final *F*_V_/*F*_M_ (**Table S5**). Importantly, leaf integrity and all its interactions were *P >* 0.05, demonstrating cut or whole leaves had negligibly different final *F*_V_/*F*_M_ (**Fig. 5; Table S5**).

**Figure 5.**
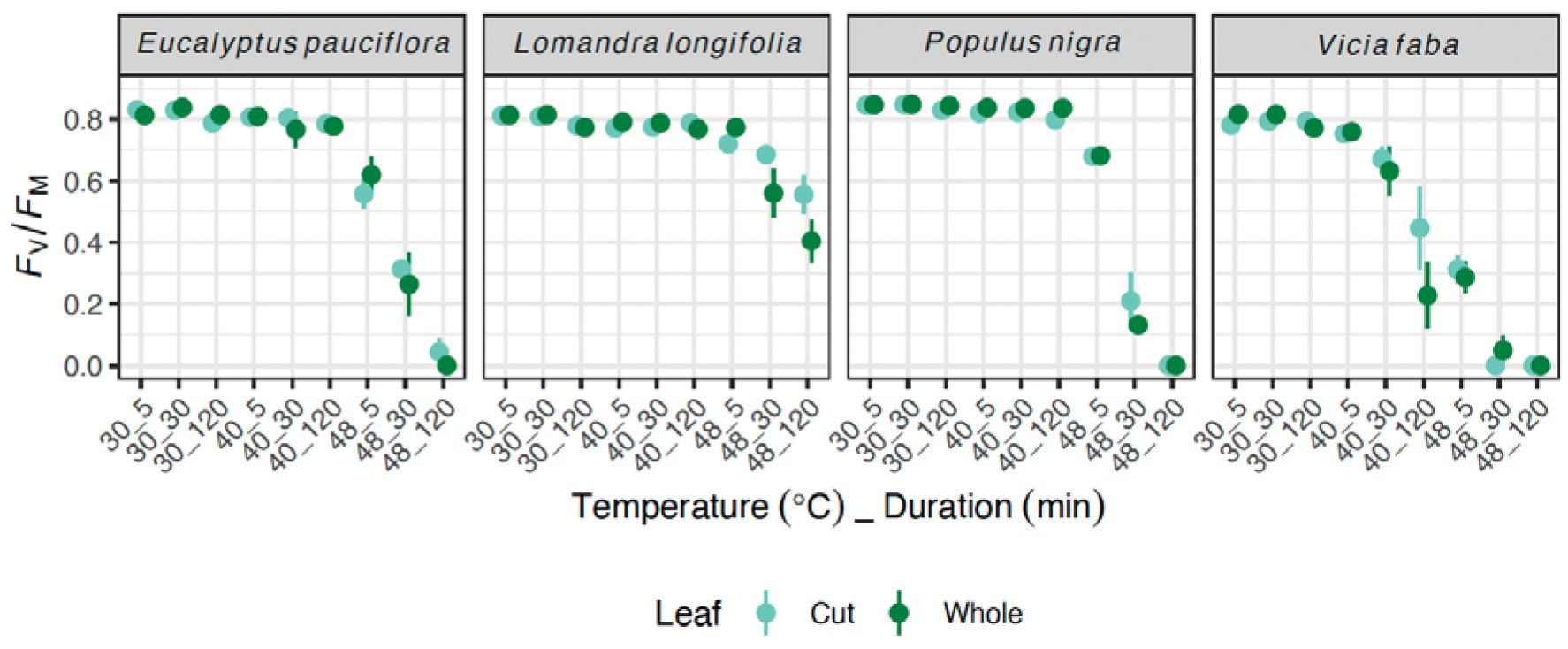
Experiment 3 findings: effects of cutting leaves on final *F*_V_/*F*_M_. Species are separated by facets, while temperature intensity and exposure duration treatments are shown together on the x-axis. Colours separate the leaf integrity (cut vs whole). Points and error bars are means ± SE.

### Experiment 4: Leaves can be collected up to 24 h prior to heat tolerance assays without compromising measurements

When collecting leaf tissue in the field there is often a delay between collection and assays. We found that time since collection had a negligible effect on *F*_V_/*F*_M_ (never significant as main effect or interaction), irrespective of initial or final measurement or the temperature × duration combination considered (**Fig. 6; Table S6**).

**Figure 6.**
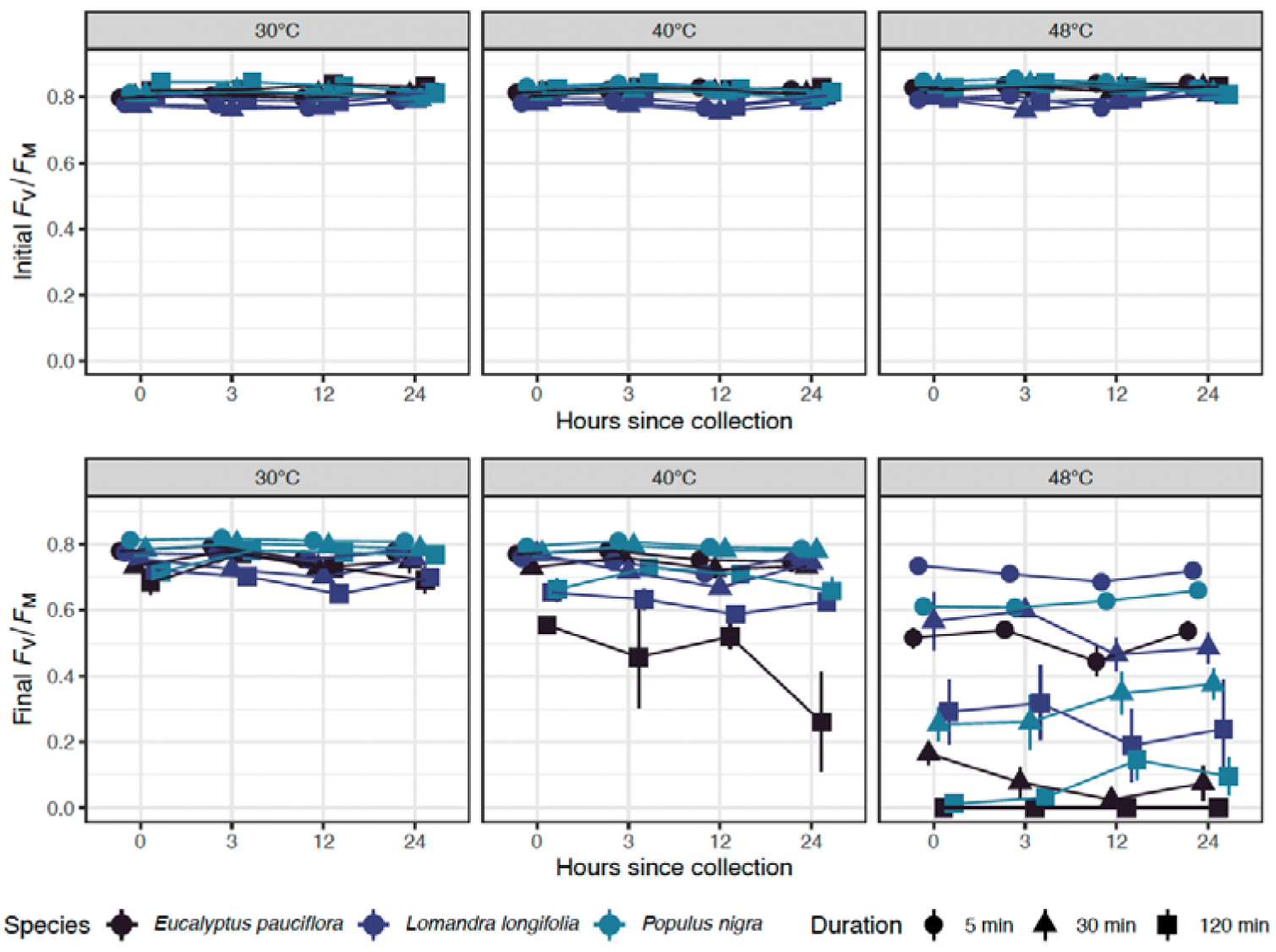
Experiment 4 findings: effects of time since collecting leaf samples on *F*_V_/*F*_M_ before (top, initial) and after (bottom, final) heat load treatments. Panels show the temperature intensity treatment, symbols separate the exposure duration treatment, and colours separate the species. Points and error bars are means ± SE.

### Experiment 5: Evidence for repair of F_V_/F_M_ over time post-heat treatment

We measured *F*_V_/*F*_M_ on the same leaf samples at multiple time points up to 24 h following heat load treatments to determine whether, after an acute heat stress, leaf tissue accumulates damage or can repair damage to restore function of *F*_V_/*F*_M_. Short durations and/or mild temperatures (e.g. 30_30, 40_5) did not lead to loss of function, and timing of measurement did not alter that result (**Fig. 7**). More intense and/or longer exposure duration (e.g. 40_120, 48_30) resulted in steep decline in *F*_V_/*F*_M_ from initial to 30 mins since treatment, and there was often an increase in *F*_V_/*F*_M_ by 7 h since treatment (**Fig. 7**). There was not a clear, ubiquitous threshold beyond which repair did not occur: *F*_V_/*F*_M_ in *L. longifolia* declined to just 0.13 at 30 mins but increased to 0.33 by 24 h since treatment (**Fig. 7**). Pairwise post-hoc tests demonstrated that the starkest change in *F*_V_/*F*_M_ from 30 mins to 24 h was consistent across species in the 40°C treatments for 30 min and 120 min exposure duration (**Table S7; Fig. 7**). Other comparisons significantly differed among species, where *E. pauciflora* and *P. nigra* were unable to recover lost function or *F*_V_/*F*_M_ further declined in 48_30 and 48_120 (**Table S7; Fig. 7**). Comparing 48_30 between *L. longifolia* and *P. nigra* reveals species-specific capability for restoration of function (**Fig. 7**). Both these species *F*_V_/*F*_M_ declines to ∼0.3 at 30 mins, but *L. longifolia* repairs and stabilises at 0.54 by 24 h while *P. nigra* initially increases to 7 h then declines to 0.22 by 24 h since treatment (**Fig. 7**). Overall, measurements taken 16–24 h are generally stable, allowing time for either repair processes to offset damage post-heat treatment or for damage to coalesce.

**Figure 7.**
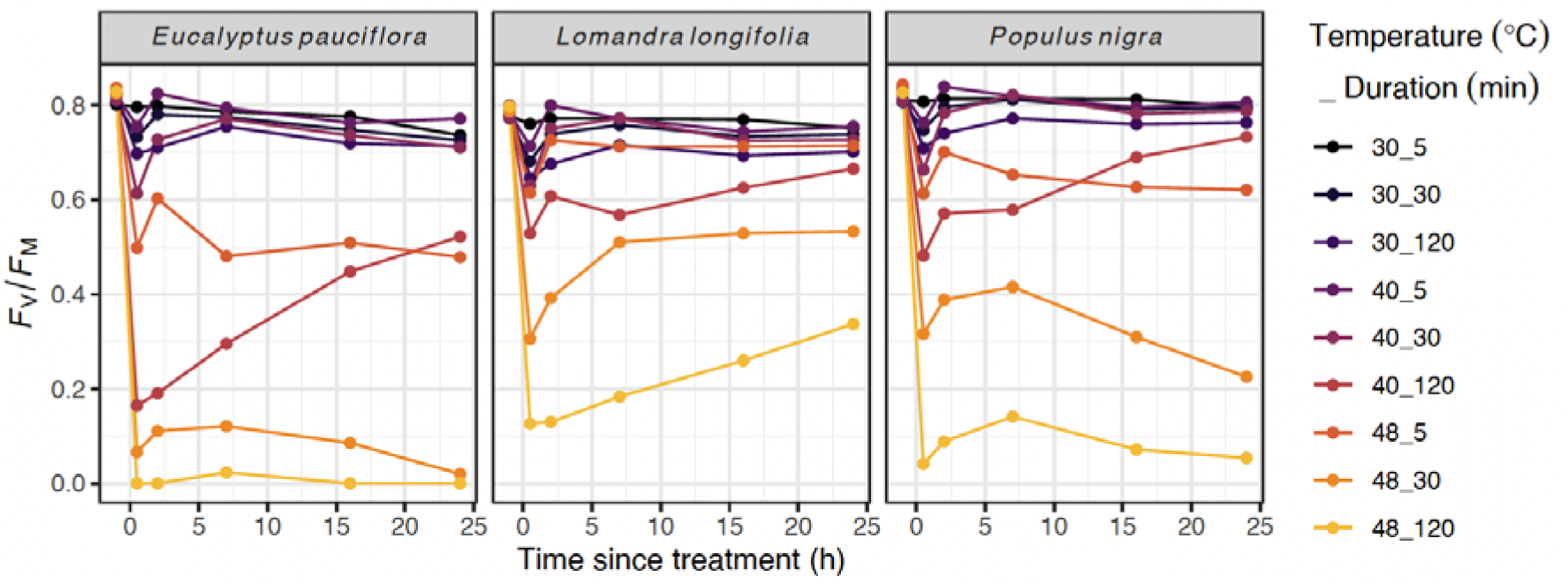
Experiment 5 findings: effects of time since heat treatment on *F*_V_/*F*_M_. Species are presented on separate panels. Colours separate the heat load (temperature intensity × exposure duration combinations).

## Discussion

Until quite recently, investigations into plant responses to heat stress have often focused on specific thermal thresholds without incorporating the cumulative nature of heat stress on organisms. As the importance of the interaction between heat stress intensity and duration for plants is becoming abundantly clear (Hüve *et al*. 2011; Teskey *et al*. 2015; Neuner & Buchner 2023; Cook *et al*. 2024; Faber, Ørsted & Ehlers 2024; Arnold *et al*. 2025a; Aitken *et al*. 2026; Noble *et al*. 2026), we developed a standard protocol to assess the thermal load sensitivity (TLS) of PSII. Our aim was to provide a robust ecophysiological approach for assessing cumulative thermal load impacts on plants that would enable comparison across regions and taxa. We tested our method across four contrasting plant forms to characterise how light conditions, leaf integrity, and time since collection influenced *F*_V_/*F*_M_ responses to heat stress. Here we have demonstrated that this approach captures the impacts of temperature intensity and exposure duration on plant functional impairment.

Based on our five experiments we advocate that the following default implementations should be suitable for interspecific comparisons of TLS using *F*_V_/*F*_M_. 1) Heat stress should be applied under sub-saturating light; 2) following heat stress the leaves should be left for 90 minutes in light; 3) use cut leaf sections to increase throughput; 4) store samples in the dark in moist conditions for up to 24 h prior to assay; and 5) a final *F*_V_/*F*_M_ measurement should be taken after 24 h. While working under dark conditions could reduce variation among studies, as the relative effect of light levels will differ among species, it will not yield biologically relevant predictors of thermal sensitivity under natural light conditions that co-occur with daytime heat stress (Krause *et al*. 2014). Below we discuss the TLS approach based on *F*_V_/*F*_M_ and identify knowledge gaps that could be addressed with this approach across diverse contexts (e.g. field, glasshouse, and controlled environment).

### Approaches to study the impacts of thermal load on plant function

Our approach aligns closely with recent studies that have recognised the importance of exposure duration or dose-dependency of thermal tolerance responses. Neuner & Buchner (2023) discuss the critical role of heat dose on functional impairment and tissue damage. They employ both visual assessments and chlorophyll fluorescence methods to evaluate heat damage and use of logistic functions to describe dose-response relationships. Their approach is similar to the TDT approach used by Cook *et al*. (2024) and Faber, Ørsted and Ehlers (2024). For example, Cook *et al*. (2024) generated tolerance threshold estimates from logistic functions to calculate *T*_50_ values and then fit log-linear regressions across exposure durations. Faber, Ørsted and Ehlers (2024) demonstrated that thermal damage to PSII accumulates additively within certain temperature ranges. Their findings highlight distinct phases where thermal effects transition from plants able to cope with stress to then causing injury and their emphasis on cumulative thermal stress mirrors our focus on non-lethal thermal thresholds. Aitken *et al*. (2026) demonstrated that the threshold of *F*_V_/*F*_M_ = 0.3 (*T*_0.3_), *CT*_max_1m_, and *z* could be used to distinguish tolerance differences between trees across a dieback gradient to a greater extent than a typical *F*_0_*-T* derived *T*_crit_. Didion-Gency *et al*. (2025) compared multiple thresholds based on *F*_V_/*F*_M_*-T*, finding that longer heat exposure (30 vs 15 mins) consistently led to lower *F*_V_/*F*_M_ and tolerance thresholds. They also tested the integrity of leaf samples (*in vivo* attached to plant vs *ex vivo* excised from plant), finding that attached samples of Aleppo pine were far more tolerant than excised ones, but lesser differences in two other coniferous species (Didion-Gency *et al*. 2025). While here we did not test attached leaves of our plants, their study highlights the need to consider the confounding effect of excising tissue on heat tolerance estimates. In our experience, maintaining hydration is key when working with excised leaf tissue for these assays – keeping samples in sealed bags with a source of moisture is generally sufficient to keep them hydrated. Minimum leaf section size (∼1 cm^2^) should be chosen to avoid ‘edge effects’ of tissue damage for *F*_V_/*F*_M_ measurements. We also note that the propensity for dehydration of target species should be considered within the timeframe of handling excised samples.

### The role of light in modulating plant responses and sensitivity to heat

Our experiments clearly show that high-intensity light conditions during heat treatment, which is more representative of natural conditions to which leaves are exposed during heat, profoundly decreased heat tolerance (all thresholds and *CT*_max_ estimates) relative to heat in darkness. Thus, plants can withstand higher maximum temperatures without the additional stress of light. Studies exploring the effect of light on heat tolerance estimates across various species demonstrate that methodological differences can alter heat tolerance estimates in either direction (e.g. Krause *et al*. 2014; Buchner *et al*. 2015; Zhang, Deng & Hao 2019; Vilas-Boas *et al*. 2023). These fundamental sources of variation make detecting a true difference in comparative meta-analyses using heat tolerance data more difficult (Perez *et al*. 2021a; Perez *et al*. 2025). Practically, our findings corroborate a recent study on 20 tree species by Nie *et al*. (2025). They identified that heat tolerance estimates from *F*_V_/*F*_M_*-T* assays were consistently reduced by light intensity during heat treatment, where 1000 µmol m^−2^ s^−1^ reduced *T*_50_ estimates by 1–5°C relative to leaves measured in darkness. Laboratory based assays are inherently artificial but taking biological context (e.g. natural irradiance exposure during heat, sun vs shade leaves) into account is a step toward more realistic tolerance estimates (Slot *et al*. 2021; Nie *et al*. 2025).

Collectively, there is compelling evidence that the effect of irradiance levels is important when estimating heat tolerance or deriving TLS parameters. We found that the thermal sensitivity parameter, *z*, which indicates how quickly plants lose heat tolerance with increasing exposure duration, is generally lower (less negative) under light conditions compared to darkness. This finding suggests that PSII of these plants is less sensitive to prolonged heat exposure (i.e. more dependent on temperature intensity than exposure duration) when also exposed to light, potentially due to the activation of photoprotective mechanisms and heat shock proteins that help mitigate thermal damage over time (Allakhverdiev *et al*. 2008). Under higher light intensity, maximum quantum yield of PSII is reduced and non-photochemical quenching mechanisms are altered. These assist with the dissipation of excessive light energy but can also be temperature dependent (Herdean *et al*. 2023), while the capacity for thermal dissipation can also be overwhelmed, leading to reduced heat tolerance (Maxwell and Johnson, 2000; Baker, 2008). Empirical studies to elucidate the dynamic interplay between light-induced photoprotection and photoinhibition, and the sources of species-specific responses to light during heat are certainly warranted.

### Exciting questions and future directions

#### 1. How do we assess the interplay of damage accumulation vs repair to restore function of heat stressed tissue?

We found enticing evidence that *F*_V_/*F*_M_ can be partially restored after an initial decline following heat exposure, and this technique expands the possibility to explore the dynamics of damage and recovery. Assessing PSII thermal sensitivity using the protocol demonstrated here will enhance the reproducibility of thermal tolerance assessments and facilitate comparative analyses across diverse plant taxa and ecological settings. Theoretical models show the potential for simplified damage and repair processes to be modelled and integrated into probabilistic frameworks for individual and population outcomes in the context of thermal stress (Ørsted, Jørgensen & Overgaard 2022; Arnold *et al*. 2025a; Noble *et al*. 2026). Yet, practical tools to resolve integrated damage-repair processes in tractable biological systems at a phenotypic level are currently limited.

The capacity for *F*_V_/*F*_M_ to be restored over time following heat exposure has recently been shown by Didion-Gency *et al*. (2025). Over four days post-heat treatment, they followed the recovery of *F*_V_/*F*_M_, *A*_sat_, and *g*_s_ from treatments between 30–60°C in three species, finding that recovery of *F*_V_/*F*_M_ to pre-heat treatment levels was not possible when temperatures exceeded 45–50°C after either 15 or 30 mins of heat exposure (Didion-Gency *et al*. 2025). Based on the consistent time-dependence of heat tolerance observed here and in other studies (Cook *et al*. 2024; Faber, Ørsted & Ehlers 2024), we would predict that longer exposure to heat treatments would further reduce the temperature beyond which *F*_V_/*F*_M_ fails to recover. Longer term observations of recovery of *F*_V_/*F*_M_ function to 14 days post-heat treatment has been undertaken by Winter *et al*. (2025). Their study on two tropical tree species found that visible tissue damage (necrosis) of leaf discs exposed to 15 min heat treatments was more closely aligned with the *T*_50_ calculated from 14 days post-heat treatment *F*_V_/*F*_M_ values (Winter *et al*. 2025). These findings demonstrate that the assumption that *T*_50_ (as typically measured) indicates irreversible loss of function does not necessarily hold. The link between decline in *F*_V_/*F*_M_ and truly irreversible damage remains poorly characterised among species. Even if the decline in *F*_V_/*F*_M_ is partially reversible (in the sense that *F*_V_/*F*_M_ values improve following heat stress), returning to pre-stress function through biosynthesis and reparation may be costly. What conditions determine whether repairing substantial heat damage to a photosynthetic module is more cost-effective than considering it ‘uneconomical to repair’ and therefore jettisoning it in favour of initiating growing a new module in its place?

#### 2. How does ecological strategy link to heat tolerance acclimation?

Among species, inherent thermal tolerance clearly differs, but pinpointing the drivers of this variation remains challenging, with climate, phylogeny, and traits explaining relatively little variance (Slot *et al*. 2021; Bison & Michaletz 2024; Briceño *et al*. 2025). However, there are several well-known drivers of PSII heat tolerance acclimation (Posch *et al*. 2025). Direct leaf temperature rather than air temperature is a significant driver of heat tolerance acclimation (Cook *et al*. 2021; Perez & Feeley 2021). While different plant species employ diverse strategies to cope with heat, leaves can generally passively avoid heat stress through anatomical structures, inclination, and transpirational cooling to reduce their temperature (Ball, Cowan & Farquhar 1988; Leigh *et al*. 2012; Leigh *et al*. 2017; Drake *et al*. 2018; Deva *et al*. 2020; Tserej & Feeley 2021). Species from contrasting origin biomes exhibit strong differences in their capacity to avoid overheating in common conditions (Arnold *et al*. 2025b). In common garden experiments, current growth conditions appear to drive PSII heat tolerance differences even among species from diverse origins (Andrew *et al*. 2023) or from contrasting parental growth conditions (Arnold *et al*. 2024).

Our four test species are vastly different in growth form, life history, and leaf structure, and their differences in thresholds and *CT*_max_ reflect that substantial variation, while *z* is somewhat less divergent. The potential for acclimation or multiple stressors to alter both *CT*_max_ and *z* in plants remains a frontier that we advocate exploring further (Arnold *et al*. 2025a; Aitken *et al*. 2026). The leaf temperature a few days prior to PSII heat tolerance measurement can be crucial for determining *T*_crit_ derived from *F*_0_-*T* (Coast *et al*. 2022; Posch *et al*. 2025; Pottinger *et al*. 2025; Hanley *et al*. 2026). Within species, thermal tolerance can acclimate, especially to high temperature (Andrew *et al*. 2023; Harris *et al*. 2024; Alvarez *et al*. 2025). One might expect a trade-off between inherent heat tolerance and acclimation capacity, but heat avoidance or heat dissipation capacity also play a role (Bison & Michaletz 2024; Arnold *et al*. 2025b). Manzi *et al*. (2025) argue that species with traits that predispose them to have higher leaf temperatures are often those that have higher heat tolerance. Both their multi-species study in Rwanda (Manzi *et al*. 2025) and another in Australia (Zhu *et al*. 2018) found that heat tolerance generally acclimated by ∼0.3°C per 1°C of growth temperature. Insight into the limits of acclimation may arise from further study on trade-offs between heat tolerance, optimal assimilation temperatures, and thermal performance breadth (Sastry, Guha & Barua 2017; Perez *et al*. 2021b).

#### 3. Can a similar approach work well for cold tolerance and help to evaluate thermal tolerance breadth?

Tolerance against chilling and extreme cold in plants remains an important aspect of abiotic stress resilience in plants (Geange *et al*. 2021). For example, climate warming can paradoxically expose plants to more extreme low temperatures and frost if snow falls as rain. There is already a theoretical foundation for cold tolerance in the TDT framework (Rezende, Castañeda & Santos 2014) and for chlorophyll fluorometry to be effective for determining cold threshold and limits for PSII (e.g. Arnold *et al*. 2021; Faber, Ørsted & Ehlers 2024; Nie *et al*. 2025). A similar exercise as we have done here for heat to determine the moderating experimental and ecological factors that affect cold tolerance thresholds and derived metrics is warranted, as there are key contextual and biophysical differences between heat and cold responses. If successful, then thermal tolerance breadth (TTB) could be comprehensively assessed with the same tools and analytical pipeline. TTB can offer insights into the fundamental limits for function at both extremes, year-round climate suitability, and adaptive capacity (Geange *et al*. 2021; Sklenář *et al*. 2023; Briceño *et al*. 2025).

## Conclusions

Standardised protocols for assessing responses of thermal load sensitivity in plants will be key to identify what ‘dose makes the poison’ (Neuner & Buchner 2023) and to ‘find the right limit’, which has been a long-debated and continually refined research field for heat tolerance studies of ectothermic animals (Clusella-Trullas *et al*. 2021; Ørsted, Jørgensen & Overgaard 2022). Our TLS protocol and analytical pipeline for *F*_V_/*F*_M_ enhance the comparability of thermal tolerance assessments across species and ecological contexts. Extensions to this approach that allow different tissue types to be directly compared (e.g. electrolyte leakage for floral and vegetative tissues, respiration for seeds and leaves) or other traits to be quantified similarly (Faber, Ørsted & Ehlers 2024) will facilitate the challenge of scaling up to whole plants (Arnold *et al*. 2025a). We hope that this approach will mature and contribute to a better understanding of the dynamics of plant resilience, acclimation, and adaptation to thermal stress. Doing so will provide valuable fundamental insights for the thermal limits for life facing a perilously hotter world, which underlies future ecological management and conservation efforts for enhancing climate resilience across systems.

## Acknowledgements

We acknowledge that this work was conducted on Ngunnawal Country and we pay our respects to the traditional custodians of these lands and elders past, present, and emerging.

## Author Contributions

P.A.A., R.J.H., S.M.A., A.M.C., A.L., and A.B.N conceptualised the study; P.A.A., R.J.H., S.M.A., and M.M.H. collected the data; P.A.A., R.J.H., S.M.A., and A.M.C. analysed the data; P.A.A. and R.J.H. wrote the manuscript; all authors contributed to editing.

## Data Availability

Data and R code for analyses and figures will be made available via Figshare and Zenodo.

