## Supplementary material for "Towards a standard approach to investigating the Thermal Load Sensitivity of photosystem II via chlorophyll fluorescence": Table S1

**Table S1:** Experiment 1 (light vs dark recovery period following heat treatment) ANOVA table: tolerance thresholds.
**Table S2:** Experiment 1 (light vs dark recovery period following heat treatment) ANOVA table: derived TLS metrics.
**Table S3:** Experiment 2 (light levels during heat treatment) ANOVA table: tolerance thresholds.
**Table S4:** Experiment 2 (light levels during heat treatment) ANOVA table: derived TLS metrics.
**Table S5:** Experiment 3 (Comparison of whole leaves vs leaf sections) ANOVA table for $F_V/F_M$.
**Table S6:** Experiment 4 (Time since leaf collection) ANOVA table for $F_V/F_M$.
**Table S7:** Experiment 5 (Time since treatment) post-hoc tests for change in $F_V/F_M$.

**Table S1:** Experiment 1 (light vs dark recovery period following heat treatment) ANOVA table for the three heat tolerance thresholds derived from the proportional change in $F_V/F_M$. Bold *P*-values indicates significance at $P <$ 0.05.

| **Experiment 1: light *vs* dark following heat treatment** | $T_{10}$ | | $T_{0.3}$ | | $T_{50}$ | | $T_{90}$ | |
|---|---|---|---|---|---|---|---|---|
| ***Factor*** | ***F*** | ***P*** | ***F*** | ***P*** | ***F*** | ***P*** | ***F*** | ***P*** |
| Light | 13.98 | **<0.001** | 33.66 | **<0.001** | 26.25 | **<0.001** | 17.80 | **<0.001** |
| Duration | 1507.88 | **<0.001** | 342.11 | **<0.001** | 735.36 | **<0.001** | 170.86 | **<0.001** |
| Species | 70.85 | **<0.001** | 184.92 | **<0.001** | 170.94 | **<0.001** | 167.84 | **<0.001** |
| Light × duration | 3.92 | **0.049** | 22.26 | **<0.001** | 9.14 | **0.003** | 25.23 | **<0.001** |
| Light × species | 2.39 | 0.076 | 2.26 | 0.091 | 1.34 | 0.267 | 3.94 | **0.012** |
| Duration × species | 11.87 | **<0.001** | 19.08 | **<0.001** | 15.35 | **<0.001** | 39.78 | **<0.001** |
| Light × duration × species | 1.25 | 0.292 | 1.27 | 0.288 | 0.26 | 0.857 | 2.16 | 0.095 |

**Table S2:** Experiment 1 (light vs dark recovery period following heat treatment) ANOVA table for the derived TLS metrics: $CT_{max\_1m}$, $CT_{max\_1h}$, and *z*. Bold *P*-values indicates significance at $P < 0.05$.

| **Experiment 1: light *vs* dark following heat treatment** | $CT_{max\_1m}$ | | $CT_{max\_1h}$ | | *z* | |
|---|---|---|---|---|---|---|
| ***Factor*** | ***F*** | ***P*** | ***F*** | ***P*** | ***F*** | ***P*** |
| Light | 28.04 | **<0.001** | 10.65 | **0.002** | 9.31 | **0.004** |
| Threshold ($T_{10}$, $T_{0.3}$, $T_{50}$, $T_{90}$) | 318.15 | **<0.001** | 3121.72 | **<0.001** | 85.39 | **<0.001** |
| Species | 42.22 | **<0.001** | 316.97 | **<0.001** | 10.35 | **<0.001** |
| Light × threshold | 11.02 | **<0.001** | 4.83 | **0.003** | 11.16 | **<0.001** |
| Light × species | 1.57 | 0.211 | 0.72 | 0.549 | 0.66 | 0.584 |
| Threshold × species | 3.07 | **0.002** | 100.68 | **<0.001** | 19.96 | **<0.001** |
| Light × threshold × species | 4.17 | **<0.001** | 1.86 | 0.065 | 2.40 | **0.016** |

**Table S3:** Experiment 2 (light levels during heat treatment) ANOVA table for the three heat tolerance thresholds derived from the proportional change in $F_V/F_M$. Bold *P*-values indicates significance at $P < 0.05$.

| **Experiment 2: light levels during heat treatment** | $T_{10}$ | | $T_{0.3}$ | | $T_{50}$ | | $T_{90}$ | |
|---|---|---|---|---|---|---|---|---|
| ***Factor*** | ***F*** | ***P*** | ***F*** | ***P*** | ***F*** | ***P*** | ***F*** | ***P*** |
| Light | 103.94 | **<0.001** | 139.31 | **<0.001** | 71.05 | **<0.001** | 83.09 | **<0.001** |
| Duration | 671.61 | **<0.001** | 341.63 | **<0.001** | 389.58 | **<0.001** | 332.42 | **<0.001** |
| Species | 295.28 | **<0.001** | 292.43 | **<0.001** | 156.46 | **<0.001** | 186.58 | **<0.001** |
| Light × duration | 2.54 | 0.057 | 2.99 | **0.032** | 1.69 | 0.170 | 5.48 | **0.001** |
| Light × species | 4.15 | **<0.001** | 2.83 | **0.004** | 2.36 | **0.015** | 2.18 | **0.024** |
| Duration × species | 2.08 | 0.103 | 20.18 | **<0.001** | 6.20 | **<0.001** | 14.15 | **<0.001** |
| Light × duration × species | 3.46 | **<0.001** | 0.90 | 0.529 | 1.41 | 0.184 | 0.97 | 0.467 |

**Table S4:** Experiment 2 (light levels during heat treatment) ANOVA table for the derived TLS metrics: $CT_{max\_1m}$, $CT_{max\_1h}$, and *z*. Bold *P*-values indicates significance at $P < 0.05$.

| **Experiment 2: light levels during heat treatment** | $CT_{max\_1m}$ | | $CT_{max\_1h}$ | | *z* | |
|---|---|---|---|---|---|---|
| ***Factor*** | ***F*** | ***P*** | ***F*** | ***P*** | ***F*** | ***P*** |
| Light | 68.30 | **<0.001** | 143.92 | **<0.001** | 10.03 | **<0.001** |
| Threshold ($T_{10}$, $T_{0.3}$, $T_{50}$, $T_{90}$) | 710.68 | **<0.001** | 2360.03 | **<0.001** | 17.88 | **<0.001** |
| Species | 96.00 | **<0.001** | 521.21 | **<0.001** | 3.96 | **0.011** |
| Light × threshold | 9.22 | **<0.001** | 16.28 | **<0.001** | 6.91 | **<0.001** |
| Light × species | 1.13 | 0.350 | 4.21 | **<0.001** | 1.05 | 0.408 |
| Threshold × species | 7.51 | **<0.001** | 34.86 | **<0.001** | 5.74 | **<0.001** |
| Light × threshold × species | 5.09 | **<0.001** | 4.70 | **<0.001** | 4.10 | **<0.001** |

**Table S5:** Experiment 3 (Comparison of whole leaves vs leaf sections) ANOVA table for $F_V/F_M$ for a subset of temperature and duration combinations. Bold *P*-values indicates significance at $P < 0.05$.

| **Experiment 3: Comparison of whole leaves vs leaf sections** | $F_V/F_M$ | |
|---|---|---|
| ***Factor*** | ***F*** | ***P*** |
| Leaf integrity | 1.51 | 0.222 |
| Temperature _ duration | 590.04 | **<0.001** |
| Species | 145.22 | **<0.001** |
| Leaf × temperature _ duration | 1.82 | 0.075 |
| Leaf × species | 0.339 | 0.797 |
| Temperature _ duration × species | 33.24 | **<0.001** |
| Leaf × Temperature _ duration × species | 1.38 | 0.119 |

**Table S6:** Experiment 4 (Time since leaf collection) ANOVA table for $F_V/F_M$ for a subset of temperature and duration combinations. Bold *P*-values indicates significance at $P < 0.05$.

| **Experiment 4: Time since leaf collection** | $F_V/F_M$ | |
|---|---|---|
| ***Factor*** | ***F*** | ***P*** |
| Time since collection | 0.39 | 0.759 |
| Measurement (initial, final) | 406.06 | **<0.001** |
| Temperature _ duration | 55.30 | **<0.001** |
| Species | 8.25 | **<0.001** |
| Time since collection × measurement | 0.17 | 0.919 |
| Time since collection × Temperature _ duration | 0.11 | 1.000 |
| Measurement × Temperature _ duration | 60.38 | **<0.001** |
| Time since collection × measurement × Temperature _ duration | 0.11 | 1.000 |

**Table S7:** Experiment 5 (Time since treatment) post-hoc tests for change in $F_V/F_M$ measured as 30 min to 24 h post-heat treatment for a subset of treatment (temperature _ duration) combinations. Bold *P*-values indicates significance at $P < 0.05$.

| **Species** | **Treatment** | **Estimate** | ***t*-ratio** | ***P*-value** |
|---|---|---|---|---|
| *Eucalyptus pauciflora* | 30_5 | 0.06 | 2.35 | **0.019** |
| | 30_30 | 0.007 | 0.26 | 0.796 |
| | 30_120 | -0.016 | -0.63 | 0.530 |
| | 40_5 | -0.016 | -0.64 | 0.523 |
| | 40_30 | -0.097 | -3.82 | **<0.001** |
| | 40_120 | -0.357 | -14.08 | **<0.001** |
| | 48_5 | 0.020 | 0.79 | 0.430 |
| | 48_30 | 0.046 | 1.83 | 0.068 |
| | 48_120 | 0.000 | 0.00 | 1.000 |
| *Lomandra longifolia* | 30_5 | 0.008 | 0.34 | 0.738 |
| | 30_30 | -0.057 | -2.26 | **0.024** |
| | 30_120 | -0.056 | -2.21 | **0.027** |
| | 40_5 | -0.042 | -1.65 | 0.100 |
| | 40_30 | -0.097 | -3.83 | **<0.001** |
| | 40_120 | -0.137 | -5.39 | **<0.001** |
| | 48_5 | -0.100 | -3.94 | **<0.001** |
| | 48_30 | -0.227 | -8.95 | **<0.001** |
| | 48_120 | -0.210 | -8.30 | **<0.001** |
| *Populus nigra* | 30_5 | 0.012 | 0.47 | 0.641 |
| | 30_30 | -0.047 | -1.87 | 0.062 |
| | 30_120 | -0.055 | -2.16 | **0.031** |
| | 40_5 | -0.042 | -1.67 | 0.096 |
| | 40_30 | -0.125 | -4.95 | **<0.001** |
| | 40_120 | -0.252 | -9.93 | **<0.001** |
| | 48_5 | -0.008 | -0.32 | 0.751 |
| | 48_30 | 0.091 | 3.59 | **<0.001** |
| | 48_120 | -0.013 | -0.51 | 0.613 |
